# Mucosal vaccine-elicited IgA is protective against zoonotic Betacoronavirus challenge

**DOI:** 10.64898/2026.08.26.747300

**Authors:** Junghwa Seo, Emma Buck, Bryan Machani, Oscar Murillo, Breetika Maharjan, Renata Filler, Kevin O. Saunders, Craig B. Wilen, Benjamin Israelow, David R. Martinez

## Abstract

Current vaccines for respiratory viruses are primarily administered intramuscularly. Messenger RNA-lipid nanoparticle (LNP)-based intramuscular vaccination for respiratory coronaviruses induces strong systemic IgG antibody responses; they do not consistently elicit IgA in the upper and lower respiratory tracts. Using a synthetic consensus spike protein aimed to broadening immunity against SARS-like viruses, SarbConS, coupled to ferritin nanoparticles co-delivered with mastoparan-7 and an FDA-approved CpG adjuvant, we intranasally boost SARS-CoV-2-immune animals. This intranasal boosting strategy elicits durable mucosal IgA responses in the respiratory tract, along with robust systemic IgG responses, and demonstrates durable protection against SARS-CoV-2 and zoonotic SARS-like viruses from bats and pangolins. Intranasal delivery of mastoparan-7 and CpG with MERS-CoV spike protein similarly elicits MERS-CoV-specific mucosal IgA and protects against MERS-CoV challenge in mice. Moreover, we observe durable protection against these genetically divergent zoonotic SARS-like viral challenges compared to intramuscular mRNA-LNP or unadjuvanted intranasal spike boosters. Intranasal SarbConS-ferritin nanoparticle intranasal vaccination similarly elicited durable mucosal IgA and antigen-specific memory B cell responses in the airways. The protective efficacy of M7-CpG adjuvanted SarbConS ferritin nanoparticle intranasal boosters was abolished in IgA knockout mice, suggesting a requirement for IgA in mediating respiratory mucosal vaccine-mediated protection against coronavirus infection. Altogether, our results demonstrate that respiratory mucosal vaccination can elicit durable and cross-protective mucosal IgA responses against genetically diverse zoonotic coronaviruses with implications for improved mucosal vaccines for highly transmissible respiratory viral pathogens.

**Highlights:**

- M7-CpG adjuvant combination elicits durable mucosal IgA responses in the respiratory tract and intestinal mucosa.
- SarbConS ferritin nanoparticle booster adjuvanted with M7-CpG increases antigen-specific memory B cells in airway-draining lymph nodes.
- Intranasal mucosal boosting with spike protein adjuvanted with M7-CpG elicits durable mucosal IgA and protects against diverse Betacoronavirus challenge.
- Vaccine-elicited IgA is required for protection against coronavirus challenges in the respiratory tract.

## Introduction

Respiratory coronaviruses, including SARS-CoV, MERS-CoV, and SARS-CoV-2, can cause significant disease. The emergence of SARS-CoV-2 in late 2019 caused a global pandemic, coronavirus disease 2019 (COVID-19). Pandemic response in the form of vaccine development led to the successful development of several vaccines that prevent severe disease^1–3^. These SARS-CoV-2 vaccines have been widely used in people and are given only intramuscularly. Despite their high degree of protection against severe disease and hospitalization following SARS-CoV-2 infection^4^, these vaccines do not reliably or consistently elicit mucosal immunity, such as mucosal IgA^5–7^. Dimeric mucosal IgA in the respiratory tract is thought to be important in protection against mucosal viral infection. Compared to SARS-CoV-2-convalescent or unvaccinated individuals, intramuscularly vaccinated individuals who received the Pfizer or other widely used intramuscular SARS-CoV-2 vaccines did not have a difference in the replication levels of SARS-CoV-2 in the respiratory tract^8–10^, potentially suggesting poor induction of mucosal immunity. Observational studies in humans demonstrate that a sustained mucosal IgA response in the upper airway is key to mediating long-lasting protection for at least 7 months against SARS-CoV-2 infection^11^. Intramuscularly delivered SARS-CoV-2 mRNA-LNP vaccines do not reliably elicit mucosal IgA in humans^6^. Moreover, increased nasal mucosal IgA levels are significantly associated with a reduced viral burden in the upper respiratory tract against influenza virus^12^. Therefore, improved vaccination strategies are needed that can reliably and consistently elicit mucosal immunity in the respiratory tract.

FluMist, a vaccine against influenza virus, is the only intranasally administered vaccine against any respiratory viral pathogen^13^. Thus, all other vaccines against human respiratory viruses – including SARS-CoV-2, respiratory syncytial virus, measles virus, mumps virus, and others – are given intramuscularly^1,14–17^. These intramuscularly administered vaccines do not elicit consistent or durable mucosal IgA in the respiratory tract, the infection site for respiratory viruses ^6,16,18^. Therefore, current vaccination strategies and approaches need to innovate to elicit durable immunity in the respiratory mucosal tract to control viral replication and reduce transmission^14^. As a mucosal IgA response following SARS-CoV-2 infection is thought to be required for longer-lasting protection^11^, experimental respiratory mucosal vaccines will likely need to elicit durable IgA responses in the respiratory tract to ideally reduce lung virus replication and reduce transmission.

While studies have focused on eliciting tissue-resident immunity in the respiratory tract^19,20^, these studies have not evaluated the durability of the response. Therefore, we do not know if IgA and protection in the respiratory mucosa can be durably elicited. Studies in animal models have reported significant improvements in immunogenicity against pan-sarbecoviruses, including elevated serum and bronchoalveolar lavage (BAL) anti-spike IgA and IgG, robust neutralizing antibodies, protective CD8+ T cell responses, and protection against viral challenge^21–23^. However, durability of the immune response that provides protection against viral challenge beyond peak immunogenicity for existing mucosal boosting approaches is unknown. Moreover, as most of the world’s population has pre-existing SARS-CoV-2 from infection, vaccination, or both, mucosal vaccines must be tested in the context of pre-existing immunity.

Here, we demonstrate that intranasal boosting of pre-existing systemic SARS-CoV-2 immunity derived from mRNA-LNP vaccines can elicit consistent IgA responses in the mucosa and respiratory tract. Using a vaccine composed of mast cell activator mastoparan-7 (M7) and TLR-9 agonist CpG adjuvants mixed with a synthetic spike antigen conjugated to ferritin nanoparticles, we demonstrate that this intranasal booster can durably elicit IgA in mice. Moreover, we observed protection against SARS-like viral challenges and demonstrated that protection against coronavirus infection depends on IgA. This study demonstrates proof-of-concept that optimizing mucosal adjuvant combinations coupled with nanoparticle vaccines can provide durable immunity in the respiratory mucosa that is protective against beta-coronaviruses. Our results have applications beyond respiratory coronavirus infections in humans and have implications for mucosal vaccines against other respiratory viruses such as influenza virus, RSV, and pandemic influenza strains such as H5N1.

## Results

### Mucosal adjuvant selection to elicit a durable IgA response

As most of the world’s population has SARS-CoV-2 immunity from infection, vaccination, or both, vaccines that aim to boost mucosal immunity must do so in the context of pre-existing SARS-CoV-2 immunity. Therefore, we modeled in SARS-CoV-2-immune animals how intranasal boosting without and with adjuvants elicits durable IgA responses. To first select an optimal adjuvant(s), we immunized mice intramuscularly with SARS-CoV-2 mRNA-LNP and boosted them intranasally with SARS-CoV-2 spike protein without and with various adjuvants that included a mast cell agonist. As mast cells line the respiratory mucosal tract^24^, mucosal adjuvants that either alone or in combination target mast cells and other antigen-presenting cells (APCs) may be advantageous to elicit a tissue-resident and durable IgA B cell response. Mastoparan-7 (M7) is a mast cell agonist that has previously been used for influenza virus vaccination in combination with short, synthetic DNA molecules containing unmethylated cytosine-guanine oligodeoxynucleotide (CpG ODN)^25^. M7 and CpG, which form stable self-assembled nanoparticles (NPs), exhibit high efficacy in inducing nasal and fecal IgA production along with CD4^+^ T cell responses when co-administered with the influenza HA^25,26^. We therefore evaluated the synergistic effects of combining the mast cell activator M7 with various Toll-like receptor (TLR) agonists to select an optimized mucosal adjuvant that elicited high magnitude and durable levels of mucosal IgA. Mice were primed intramuscularly with SARS-CoV-2 mRNA-LNP (Comirnaty BNT162b2) and subsequently boosted intranasally at weeks three and six with SARS-CoV-2 spike (S) protein formulated with different adjuvant combinations (Fig. S1A). SARS-CoV-2 spike-specific systemic IgG and mucosal IgA levels were monitored longitudinally from week 2 to week 16, utilizing fecal samples as a non-invasive readout for mucosal immunity. Two weeks post prime immunization, minor induction of serum IgG and IgA, as well as fecal IgG, was observed, while fecal IgA was barely detectable (Fig. S1B-S1E). At week 5, 2 weeks post-1^st^ boost, both IgG and IgA induction showed a moderate increase. We therefore asked if an additional intranasal boost at week 6 increased mucosal IgA. At week 8, following a second intranasal boost, a marked increase in mucosal IgA was observed (Fig. S1B-E). Moreover, at week 12, which was 6 weeks post the 2^nd^ intranasal boost, mucosal IgA continued to increase. Based on the increasing mucosal IgA levels well after the 2^nd^ intranasal boost, we chose to implement two intranasal boosts for subsequent vaccinations. At week 8, the M7-CpG group exhibited a superior systemic IgG response compared to the group that received an intramuscular boost with SARS-CoV-2 mRNA-LNP (Fig.S1B, S1D). To assess the durability of mucosal immunity, we measured IgA levels in fecal samples and found that M7-CpG and M7-MPLA elicited the highest fecal IgA levels at week 8 compared to either the intramuscular boost or the unadjuvanted intranasal boost. Furthermore, at weeks 12 and 16, the M7-CpG group displayed the highest serum and fecal IgA titers among all experimental groups (Fig. S1C, S1E). Consequently, we used M7-CpG as the experimental mucosal adjuvant combination to evaluate vaccine-elicited immunity in the respiratory tract and to measure protection against viral challenge.

### Protein nanoparticles admixed with combination adjuvants delivered intranasally elicit durable nasal mucosa IgA responses in the upper respiratory tract

After establishing M7-CpG as an adjuvant combination that elicited the most durable and the highest magnitude of mucosal IgA, we evaluated whether an intranasal booster employing ferritin nanoparticles displaying a consensus spike antigen was similarly immunogenic. The consensus spike antigen, SarbConS, was designed by creating a consensus sequence of 23 SARS-like spikes to capture genetic and antigenic diversity among SARS-like viruses (Fig. 1A). We first confirmed that SarbConS spike protein was similarly immunogenic when delivered intranasally. We also evaluated the durability of antibody responses elicited by the M7-CpG-adjuvanted SarbConS intranasal boost; serum and fecal samples were collected longitudinally up to week 20 post-prime (Fig. S2A). Remarkably, intranasal delivery of M7-CpG adjuvanted SarbConS protein demonstrated superior mucosal IgA responses compared to the intramuscular mRNA-LNP boost (Fig. S2B-E). Importantly, no significant spike-specific IgE responses were detected in the serum (Fig. S2F, S2G) or the respiratory tract (Fig. S2H, S2I) of M7-CpG adjuvanted SarbConS ferritin nanoparticle intranasal boost group. Together, these results indicate that M7-CpG coupled with protein nanoparticle vaccine delivery via the respiratory mucosa is highly effective in eliciting durable mucosal IgA responses and systemic IgG responses.

**Figure 1.**
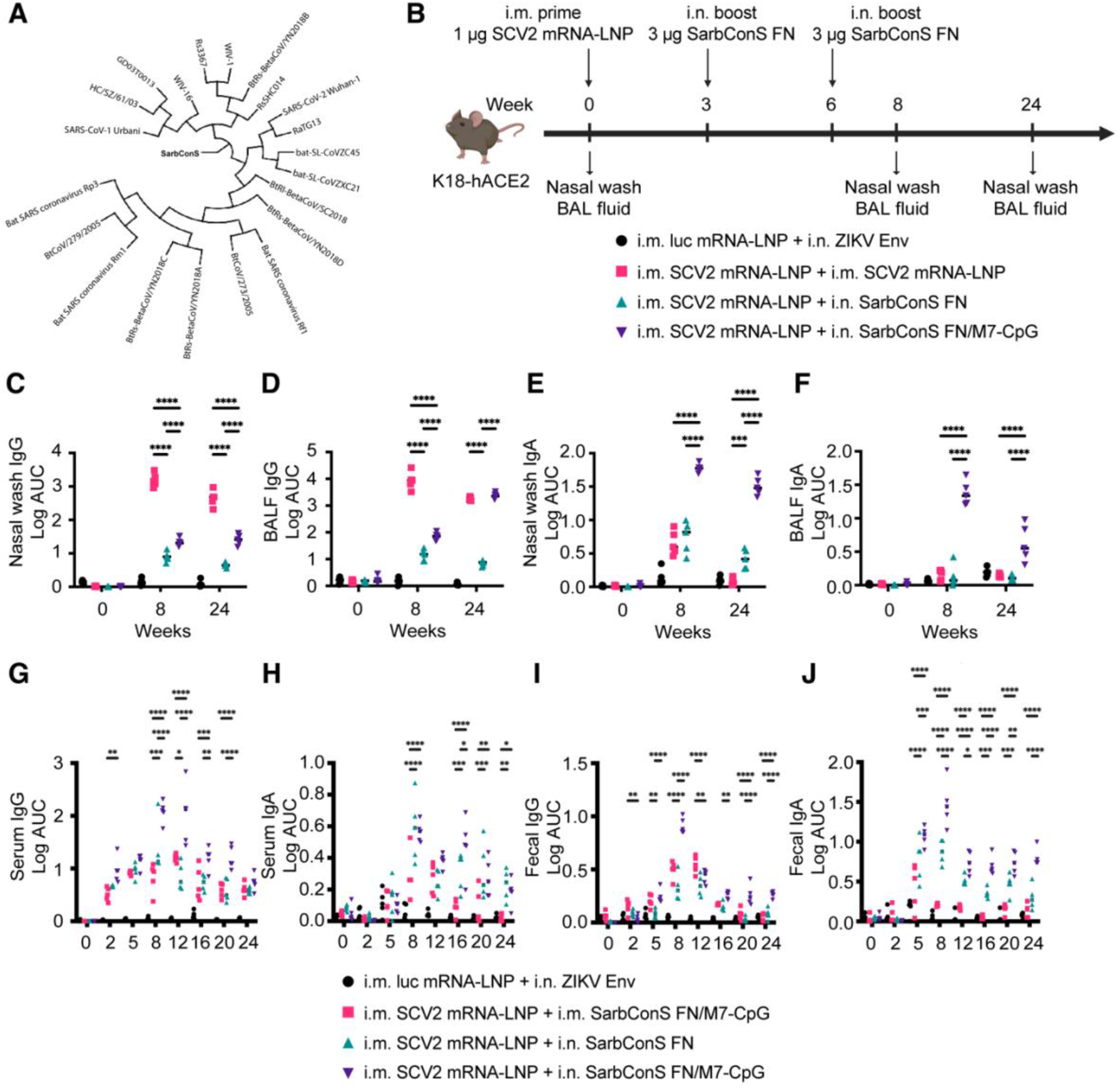
A ferritin nanoparticle intranasal booster admixed with M7-CpG elicits durable mucosal IgA and systemic IgG responses in the airway mucosa. Evaluation of spike-specific IgA and IgG responses in the mucosal and systemic compartments following intranasal boosting of SarbConS ferritin nanoparticle adjuvanted with combination adjuvant M7-CpG. K18-hACE2 mice were immunized intramuscularly at week 0 and intranasally at weeks 3 and 6. Nasal wash, bronchoalveolar lavage fluid (BALF), feces, and serum were collected at the indicated time points (*n* = 5 per group). **(A)** Phylogenetic relationship of SarbConS with SARS-like coronaviruses. **(B)** Experimental timeline for the immunization and specimen collection. **(C-F), (G-J)** Spike-specific IgA and IgG levels in each sample were measured by ELISA. Data are presented as mean ± SEM. Statistical significance was determined using two-way ANOVA followed by Tukey’s multiple comparisons test. *p < 0.05, **p < 0.01, ***p < 0.001, and ****p < 0.0001.

Ferritin nanoparticles offer intrinsic advantages as they exhibit increased structural stability and spontaneously self-assemble into 24-mer nanospheres. By displaying coronavirus spike protein in a repetitive array, ferritin nanoparticles drive robust B cell activation and broadly neutralizing antibodies when given intramuscularly^27–29^. However, it is not known if mucosal delivery of protein nanoparticles, including those presented by a ferritin scaffold, will be immunogenic and protect against viral challenge in the respiratory tract. To assess the durability of antibody responses in the respiratory airway elicited by intranasally boosting with SarbConS ferritin nanoparticles adjuvanted with M7-CpG, nasal wash and bronchoalveolar lavage fluid (BALF) at week 8 and week 24 post-prime to measure the IgA and IgG response. SarbConS-specific IgG and IgA titers were measured by ELISA. At week 24, nasal wash IgG titers were highest in the intramuscular SARS-CoV-2 mRNA-LNP prime and boost group (Fig. 1C). In contrast, BALF IgG titers at week 24 remained comparably high in intranasally boosted groups with SarbConS ferritin nanoparticles adjuvanted with M7-CpG, showing no significant difference compared to the intramuscular SARS-CoV-2 mRNA-LNP boost group (Fig. 1D). Importantly, nasal mucosal IgA was induced to high levels by the M7-CpG adjuvanted SarbConS ferritin nanoparticle intranasal boost (Fig. 1E). Similarly, the most durable IgA responses were generated in M7-CpG-adjuvanted SarbConS ferritin nanoparticle intranasal boost (Fig. 1E). BALF IgA levels were highest and remained at high levels at week 24 in the M7-CpG-adjuvanted SarbConS ferritin nanoparticle intranasal boost (Fig. 1F). Compared to other groups, the M7-CpG adjuvanted SarbConS ferritin nanoparticle intranasal boost elicited the highest levels of serum IgG, serum IgA, fecal IgG, and fecal IgA overtime and remained the most durable through 24-weeks post prime (Fig. 1G, 1H, 1I, 1J). These results suggest that intranasal delivery of M7-CpG adjuvanted ferritin nanoparticles expressing SarbConS elicit durable IgA responses in the upper respiratory tract.

### Protein nanoparticles admixed with a combination of adjuvants delivered intranasally expand memory B cell responses in the airway

As the M7-CpG-adjuvanted SarbConS ferritin nanoparticle intranasal boost elicited the highest and most durable levels of mucosal IgA, we hypothesized that antigen-specific memory B cells underwent expansion in airway-draining lymph nodes. To address this, we measured the frequency of antigen-specific memory B cells in the lungs, mediastinal lymph nodes, and cervical lymph nodes in vaccinated mice at weeks 8 and 24. At week 8, the frequency of antigen-specific (PE^+^APC^+^ double positive SarbConS) IgD^-^B220^+^CD19^+^CD38^+^ cells (Fig. S3) was significantly increased in the M7-CpG-adjuvanted SarbConS ferritin nanoparticle intranasally vaccinated group (Fig. 2A-F). Importantly, at week 24, the antigen-specific memory B cell response remained the highest, whereas no detectable memory B cell responses were observed in the unadjuvanted SarbConS ferritin nanoparticle intranasally vaccinated group (Fig. 2G-I and Fig. S4). These results suggest that combination adjuvants elicit long-lasting antigen-specific memory B cell responses in the respiratory mucosa.

**Figure 2.**
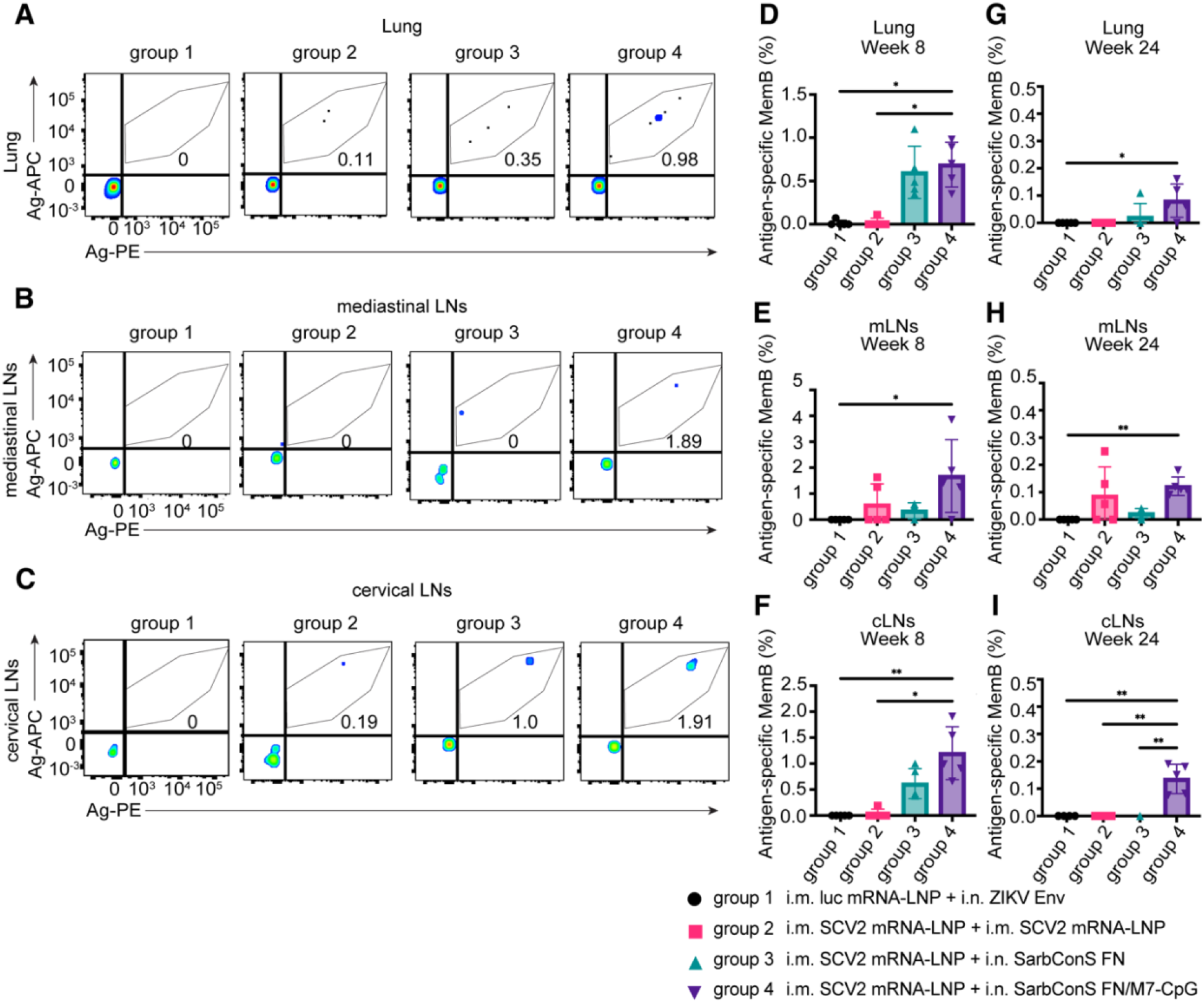
Intranasal boosting with SarbConS ferritin nanoparticle adjuvanted with M7-CpG enhances antigen-specific memory B cell responses in the lungs and airway draining lymph nodes. Flow cytometry analysis of antigen-specific memory B cell frequencies in the lungs, mediastinal lymph nodes (mLNs), and cervical lymph nodes (cLNs). **(A-C)** K18-hACE2 mice were immunized intramuscularly at week 0 and intranasally at weeks 3 and 6. Lungs and airway lymph nodes were collected and analyzed at week 8 or week 24. **(A-C)** Representative flow cytometry plots of antigen-specific CD3ε^-^IgD^-^B220^+^CD19^+^CD38^+^ memory B cells at week 8 in lungs **(A)**, mediastinal LNs **(B),** and cervical LNs **(C)**. **(D-I)** Quantification of the frequencies of antigen-specific memory B cells at week 8 (*n* = 5 per group) **(D-F)** or week 24 (*n* = 4 or 5 per group) **(G-I)**. Data are presented as mean ± SEM. Statistical significance was determined using the Kruskal-Wallis test followed by Dunn’s multiple comparisons test. *p < 0.05, **p < 0.01.

### Broadly neutralizing and cross-reactive mucosal IgA responses elicited by intranasally delivered M7-CpG-adjuvanted SarbConS-ferritin nanoparticles

We next sought to determine whether M7-CpG-adjuvanted SarbConS ferritin nanoparticle induce IgA responses that are cross-reactive against genetically diverse zoonotic SARS-like virus spikes (Fig. 3). While we observed some cross-reactive IgA responses against SARS2-like RaTG13 spike and SARS2-like Pangolin-CoV spike in mice vaccinated intramuscularly with SARS-CoV-2 mRNA-LNP (Fig. 3A, 3D), mucosal IgA responses were highest and most consistently elicited in mice that received the intranasal delivery of M7-CpG-adjuvanted SarbConS ferritin nanoparticles. Specifically, we observed robust spike-specific IgA responses against RaTG13 and BANAL236 in the nasal wash and BALF of the M7-CpG adjuvanted SarbConS ferritin nanoparticle intranasally vaccinated group (Fig. 3A-B, 3E-F). We also observed a similar magnitude of serum IgG responses in the M7-CpG adjuvanted intranasally boosted group compared to the intramuscularly boosted group (Fig. 3I-L). Conversely, serum IgA levels specific to RaTG13 and BANAL236 were further enhanced in the M7-CpG adjuvanted intranasally boosted group (Fig. 3M, 3N).

**Figure 3.**
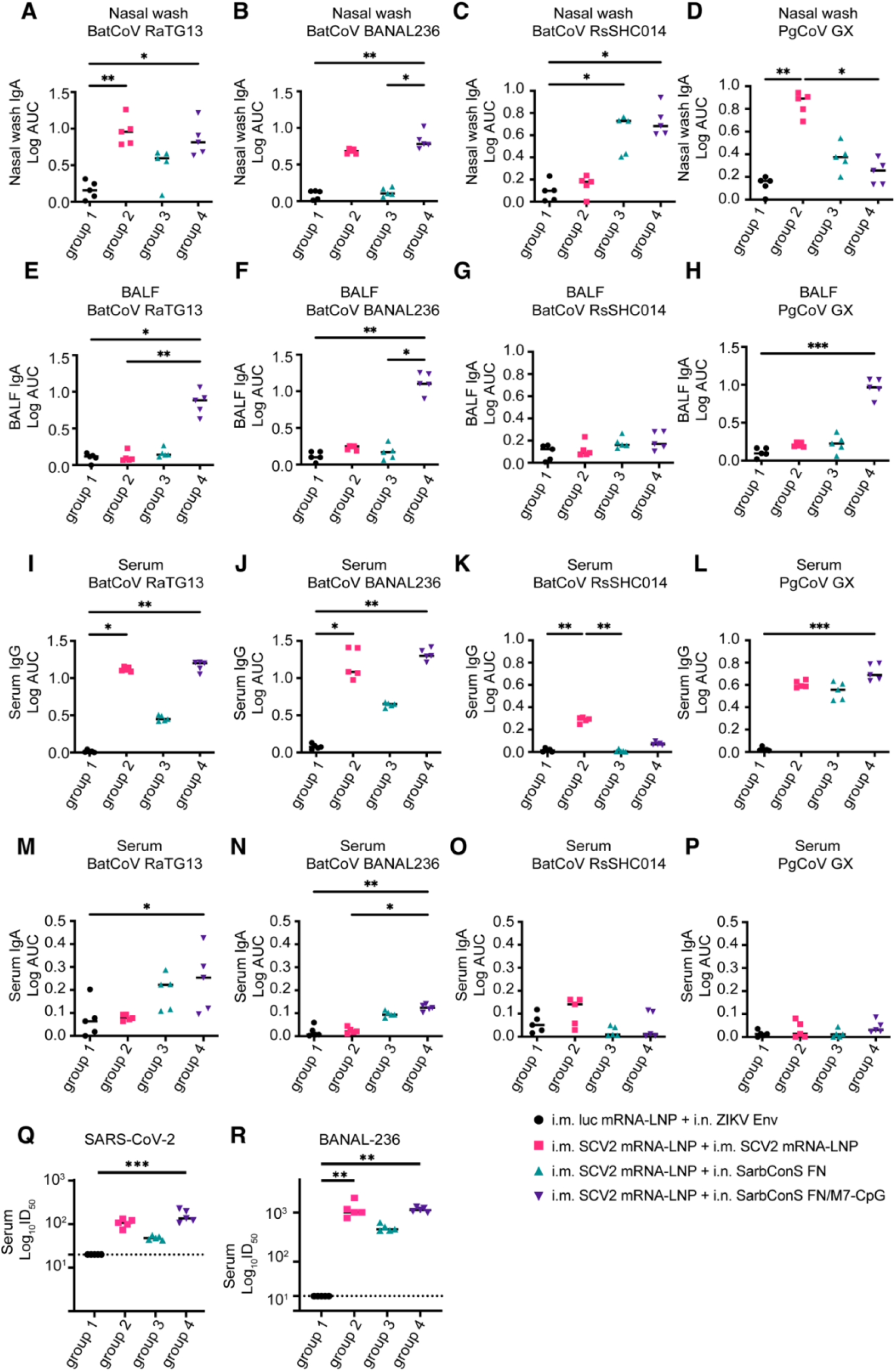
A ferritin nanoparticle intranasal booster admixed with M7-CpG produces IgA antibodies and neutralizing antibodies against zoonotic coronaviruses. Zoonotic beta-coronavirus spike-specific IgA responses in the upper and lower respiratory tract. K18-hACE2 mice were immunized as shown in Fig. 1B. At 2 weeks post 2^nd^ boost, nasal wash, BALF, and sera were harvested. Nasal mucosa (nasal wash) and BALF IgA or serum antibody titers were determined by ELISA against BatCoV RaTG13 spike **(A, E, I, M)**, BatCoV BANAL-236 S2P **(B, F, J, N)**, BatCoV RsSHC014 RBD **(C, G, K, O)**, and PgCoV-GXP4L RBD **(D, H, L, P)**. (*n* = 5 per group). Serum neutralizing antibodies were measured by pseudotyped viruses expressing SARS-CoV-2 spike **(Q)** and BANAL-236 spike **(R)**. Data are presented as mean ± SEM. Statistical significance was determined using the Kruskal-Wallis test followed by Dunn’s multiple comparisons test. *p < 0.05, **p < 0.01, and ***p < 0.001.

Finally, we measured neutralizing antibody responses in serum against SARS-CoV-2 Wuhan-1 and bat SARS2-like zoonotic BANAL-236 viruses using a pseudovirus neutralizing antibody assay. Expectedly, no neutralization was observed against SARS-CoV-2 and BANAL-236 in negative control mice. In contrast, we observed the highest serum neutralizing antibodies both against SARS-CoV-2 and BANAL-236 in mice that were intranasal boosted with M7-CpG-adjuvanted SarbConS ferritin nanoparticles (Fig. 3Q, 3R).

### Protein nanoparticles admixed with combination adjuvants delivered intranasally durably protect against zoonotic SARS-like virus challenges

We next evaluated the durability of the protective efficacy of respiratory mucosally delivered intranasal booster vaccines. In contrast to most studies that measure protective efficacy near or at peak immunogenicity, around 2 weeks following the vaccination, we assessed the durability of protection against viral challenge 10 weeks post the 2^nd^ intranasal boost. We separately challenged two cohorts of mice with high doses of SARS-like RsSHC014 and SARS2-like pangolin-CoV (Fig. 4A). By day 2 post-challenge, high levels of viral replication were present in the lung, nasal turbinate, and nasal wash for both RsSHC014 and Pangolin-CoV in the negative control group (Fig. 4B-G). In contrast, some degree of protection was afforded in the unadjuvanted SarbConS ferritin nanoparticle intranasally vaccinated group, with some reduction in the nasal turbinates against Pangolin-CoV virus replication (Fig. 4F). Remarkably, M7-CpG-adjuvanted SarbConS ferritin nanoparticle-intranasally vaccinated mice showed the most protection against both RsSHC014 and Pangolin-CoV with the most significant reduction in viral replication in the lung, nasal turbinate, and nasal wash (Fig. 4B-G). Notably, RsSHC014 and Pangolin-CoV were markedly suppressed in the nasal wash (Fig. 4D, 4G), indicating robust durable protection in the upper respiratory tract. These findings indicate that intranasal boosting with M7-CpG adjuvanted SarbConS ferritin nanoparticles effectively and durably restrict viral replication and shedding in the upper respiratory tract.

**Figure 4.**
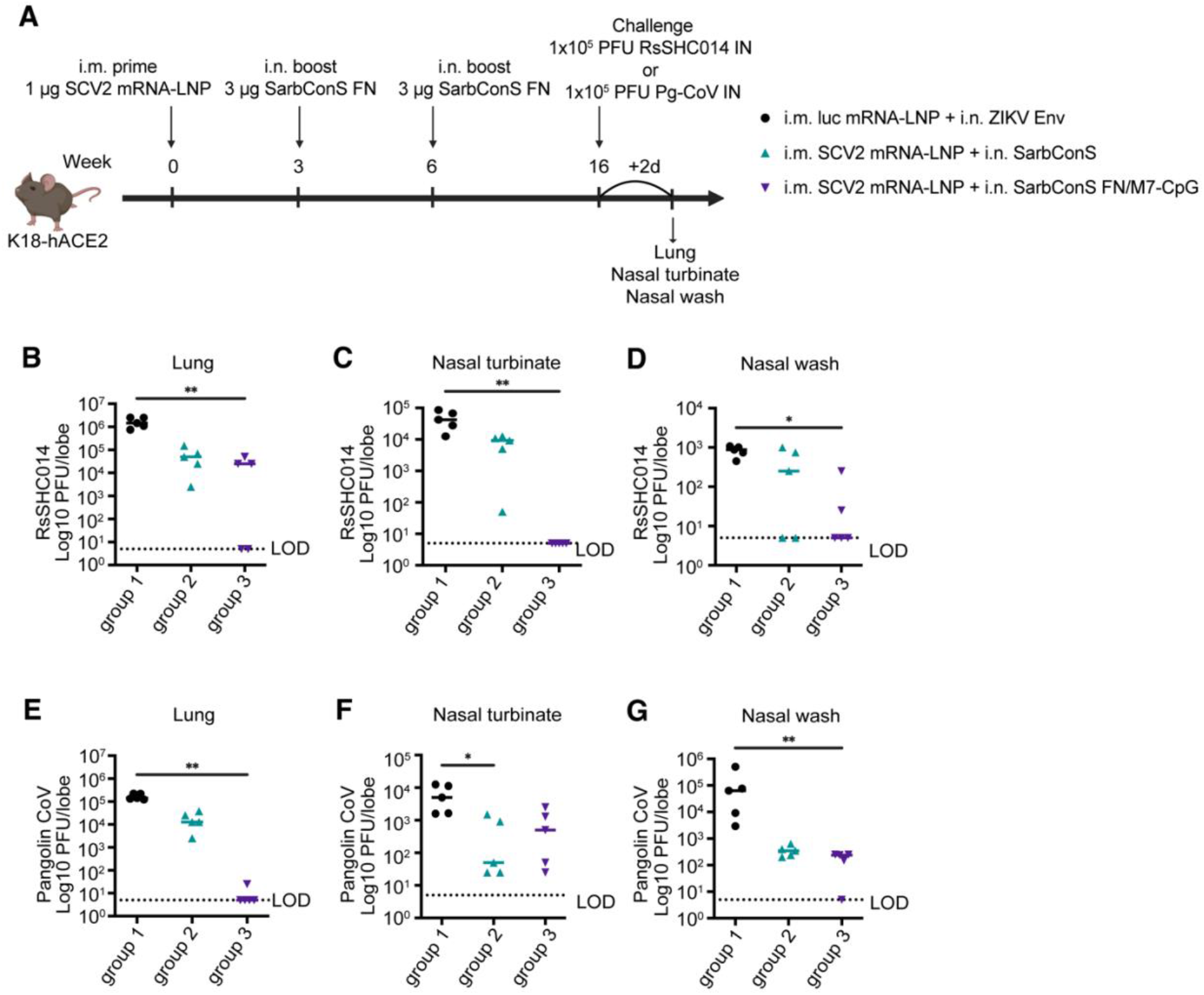
Intranasal vaccination with SarbConS ferritin nanoparticle admixed with M7-CpG confers durable cross-protection against heterologous SARS-like coronavirus challenge. Viral titers in the respiratory tract following heterologous challenges. **(A)** Experimental timeline of the vaccination and viral challenge. K18-hACE2 mice were immunized intramuscularly at week 0 and intranasally at weeks 3 and 6. At week 16, 10-weeks post 2^nd^ intranasal boost, mice were challenged with RsSHC014 (1×10^5^ PFU) or Pangolin-CoV (1×10^5^ PFU), and tissue viral titers were measured by plaque assay. **(B-G)** Mice were infected with either RsSHC014 **(B-D)** or Pangolin-CoV **(E-G)**. At 2 dpi, viral loads were measured in the lungs **(B, E)**, nasal turbinates **(C, F)**, and nasal washes **(D, G)** (*n* = 5 per group). The dashed line indicates the limit of detection (LOD), and undetectable samples were arbitrarily set to 5 PFU/tissue or PFU/mL. Data are presented as mean ± SEM. Statistical significance was determined using the Kruskal-Wallis test followed by Dunn’s multiple comparisons test. *p < 0.05, **p < 0.01.

### Ferritin NP intranasal booster provides a robust IgA response and protects against MERS-CoV challenge

To investigate whether induction of robust IgA responses by our combination-adjuvanted ferritin nanoparticle intranasal booster strategy is broadly applicable to beta-coronaviruses, we primed mice with MERS-CoV spike mRNA-LNP followed by two intranasal or intramuscular boosts with the M7-CpG-adjuvanted or unadjuvanted MERS spike ferritin nanoparticles (Fig. 5A). Vaccinated mice were challenged with the mouse-adapted MERS-CoV at two weeks post-final intranasal boost. Complete protection against lower airway viral replication in the lung was achieved across all vaccinated groups boosted either intramuscularly or intranasally compared to controls (Fig. 5B). In contrast, only the ferritin nanoparticle-intranasally boosted mice showed complete protection from replication in nasal turbinates. These results suggest that intranasal boosting is required to protect against MERS-CoV replication in the upper respiratory tract (Fig. 5C). We then measured IgG and IgA levels in both the serum and in the nasal wash and BALF. High levels of MERS-CoV spike-specific IgG titers were detected in intramuscularly or intranasally boosted M7-CpG-adjuvanted MERS-CoV spike ferritin nanoparticle boosted mice (Fig. 5D, 5F, and 5G). High levels of mucosal IgA were only observed in the intranasally boosted M7-CpG adjuvanted MERS-CoV spike ferritin nanoparticle group (Fig. 5E, 5H, and 5I). Our findings demonstrate that intranasal delivery of M7-CpG-adjuvanted MERS-CoV spike ferritin nanoparticles as a boosting strategy elicits the highest levels of mucosal IgA in the upper and lower respiratory tracts and more robustly protects against upper respiratory tract MERS-CoV infection. Thus, our boosting strategy for inducing mucosal IgA responses was applicable to multiple subgenera of beta-coronaviruses.

**Figure 5.**
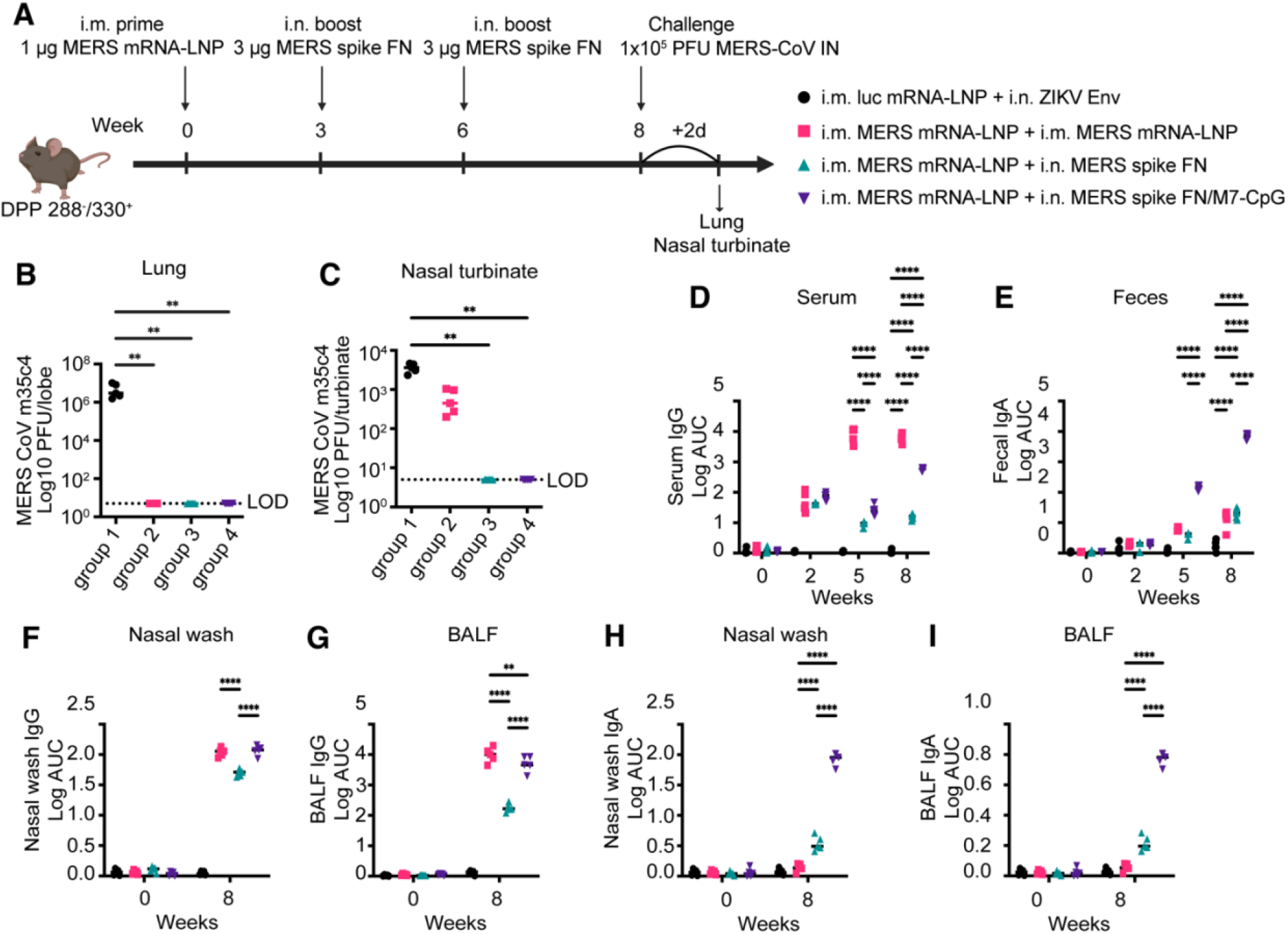
Protection efficacy of mucosal booster admixed with M7-CpG against mouse-adapted MERS-CoV (m35c4) challenge. **(A)** 6 to 12-weeks-old DPP 288^+^/330^+^ were primed with MERS mRNA-LNP intramuscularly at week 0 and boosted with adjuvanted or unadjuvanted MERS-CoV spike ferritin nanoparticle intranasally at weeks 3 and 6. At week 8, mice were challenged with MERS-CoV m35c4 at 1×10^5^ PFU. **(B, C)** Plaque assay showing viral loads in the lungs **(B)** and nasal turbinates **(C)**. The dashed line indicates the limit of detection (LOD), and undetectable samples were arbitrarily set to 5 PFU/tissue. **(D-I)** MERS-CoV spike-specific antibody titers at the indicated time points. **(D, E)** serum IgG **(D)** and fecal IgA **(E)**. **(F-I)** Respiratory tract antibody titers, showing nasal wash IgG **(F)**, BALF IgG **(G)**, nasal wash IgA **(H)** and BALF IgA **(I)**.

### IgA is required for controlling viral replication in the respiratory mucosa

We next sought to determine whether mucosal IgA was required for mediating the protective efficacy against coronavirus challenge. Like in previous intramuscular prime and intranasal boost experiments, we vaccinated IgA knockout (KO) and wild-type mice intramuscularly with SARS-CoV-2 mRNA-LNP and boosted intramuscularly vs. intranasally with M7-CpG adjuvanted SarbConS ferritin nanoparticle vaccines as well as with control vaccines (Fig. 6A). In wild-type and IgA KO mice, serum IgG levels were similar in mice vaccinated with 1) negative control vaccines, 2) SARS-CoV-2 mRNA-LNP intramuscular prime and intramuscular boost, and 3) SARS-CoV-2 mRNA-LNP intramuscular prime and SarbConS ferritin nanoparticle intranasal boost (Fig. 6B). Wild-type mice elicited high levels of mucosal IgA when vaccinated with the SARS-CoV-2 mRNA-LNP intramuscular prime and SarbConS ferritin nanoparticle intranasal boost (Fig. 6C) and, as expected, did not elicit mucosal IgA in mice vaccinated with negative control vaccines nor the SARS-CoV-2 mRNA-LNP intramuscular prime and intramuscular boost. In contrast, IgA KO mice did not elicit mucosal IgA in the SARS-CoV-2 mRNA-LNP intramuscular prime and SarbConS ferritin nanoparticle intranasal boosted groups (Fig. 6C). Similarly, negative control vaccinated mice and SARS-CoV-2 mRNA-LNP intramuscular prime and intramuscular boost did not elicit measurable mucosal IgA responses (Fig. 6C).

**Figure 6.**
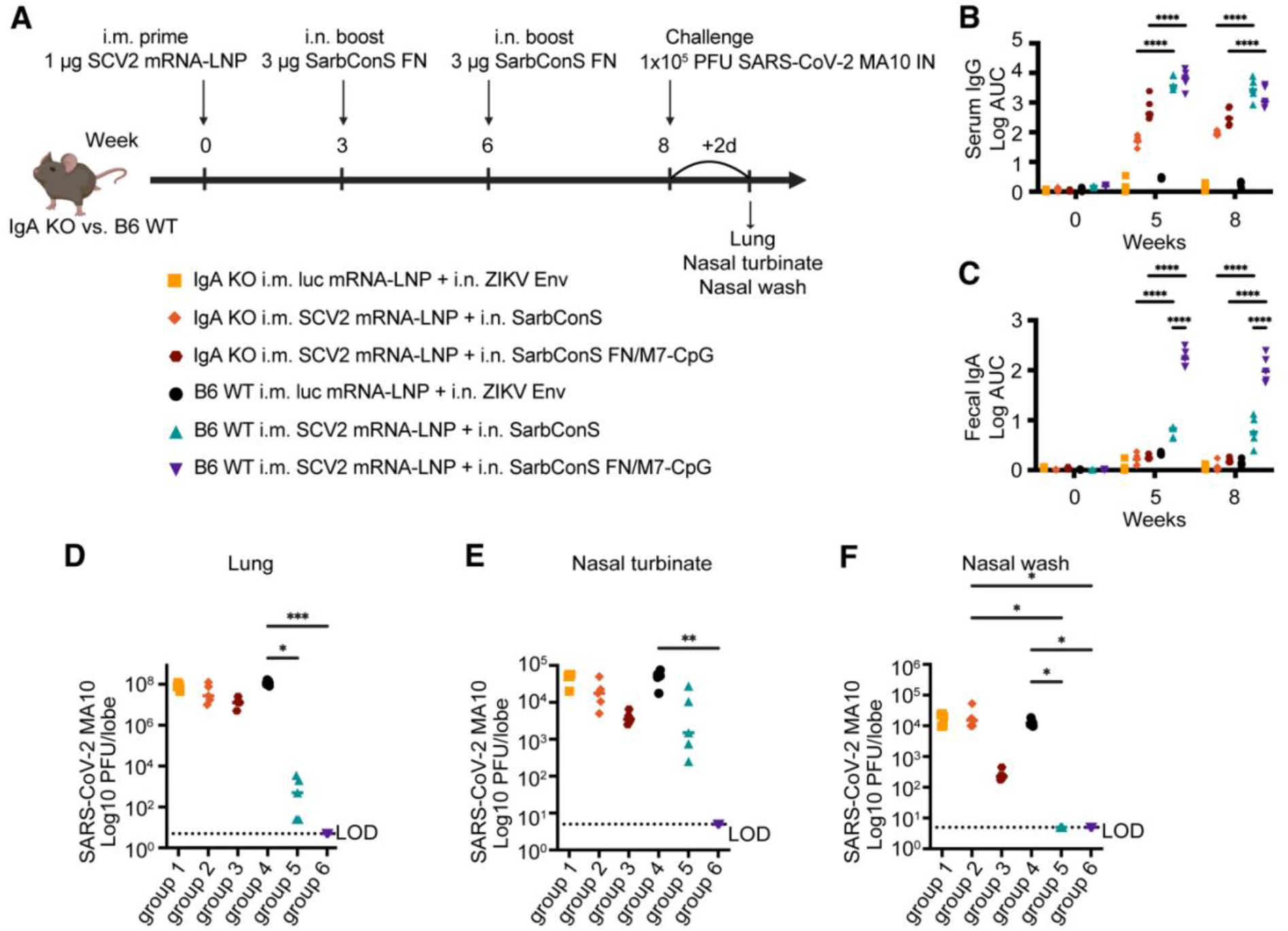
IgA is required for protection against mouse-adapted SARS-CoV-2 MA10 elicited by M7-CpG adjuvanted SarbConS ferritin nanoparticle intranasal booster. Evaluation of IgA-mediated protection against SARS-CoV-2 MA10 infection in the lower and upper respiratory tracts. **(A)** Experimental timeline of the vaccination and mouse-adapted SARS-CoV-2 (MA10) viral challenge. IgA KO or C57BL/6J wild-type mice were immunized intramuscularly at week 0 and intranasally at weeks 3 and 6. Mice were challenged with MA10 at 1×10^5^ PFU at week 8. **(B-C)** Measurement of serum IgG **(B)** and fecal IgG **(C)** titers by ELISA. **(D-F)** At 2 days post-infection (dpi), the lungs **(D)**, nasal turbinates **(E)**, and nasal washes **(F)** were collected, and viral titers were determined by plaque assay (*n* = 5 per group). Values below the limit of detection (LOD) were assigned a value of 5 PFU/tissue or PFU/mL for statistical analysis. Data are presented as mean ± SEM. Statistical significance was determined using the Kruskal-Wallis test followed by Dunn’s multiple comparisons test. *p < 0.05, **p < 0.01, ***p < 0.001, and ****p < 0.0001.

Both wild-type and IgA KO vaccinated mice were challenged with a high dose of mouse-adapted SARS-CoV-2, and virus replication was measured at 2 dpi (Fig. 6A). M7-CpG adjuvanted SarbConS ferritin nanoparticle intranasally vaccinated wild-type mice completely controlled SARS-CoV-2 replication in the lung, nasal turbinate, and nasal wash (Fig. 6D-F). In contrast, unadjuvanted SarbConS ferritin nanoparticle intranasally vaccinated wild-type mice had some breakthrough SARS-CoV-2 replication in the lungs and nasal turbinates. Wild-type mice from the negative control group expectedly had high levels of SARS-CoV-2 replication in the lung, nasal turbinate, and nasal wash. Remarkably, complete abrogation in protection against SARS-CoV-2 was observed in both M7-CpG-adjuvanted SarbConS ferritin nanoparticle intranasally vaccinated and unadjuvanted SarbConS ferritin nanoparticle intranasally vaccinated IgA KO mice (Fig. 6D-6F). In IgA KO mice, but not in wild-type mice, we observed high levels of SARS-CoV-2 replication in the lung, nasal turbinate, and nasal wash (Fig. 6D-6F). Importantly, no significant difference in SARS-CoV-2 replication was observed in negative control vaccinated groups of wild-type mice compared to IgA KO mice, indicating that the IgA KO mice are not inherently more resistant to infection. These findings suggest that mucosal IgA in the upper and lower respiratory tracts is necessary in mediating protection in M7-CpG-adjuvanted SarbConS ferritin nanoparticle-intranasally vaccinated mice.

## Discussion

In this study, we addressed a critical gap and evaluated the durability of vaccine-induced mucosal IgA responses and determined that the durable mucosal IgA response is required for vaccine-mediated protection against coronavirus infection. A successful respiratory mucosal vaccine must elicit long-lasting mucosal responses^14^. While previous studies demonstrated that mucosal boosting following intramuscular priming results in improved mucosal immunity^7,19,30^, no study has assessed the durability of mucosal vaccination nor the requirement of IgA in mediating protection. Several studies also evaluated respiratory mucosal vaccination strategies at peak immunogenicity^20,31^, but no study has rigorously evaluated durable protection or the antibody-mediated requirement for long-lasting protection. Moreover, studies have not addressed whether cross-reactive IgA responses in the mucosa can protect against distinct genetically diverse zoonotic coronaviruses.

As demonstrated by the COVID-19 pandemic, improved vaccination strategies are needed to more rapidly control respiratory coronavirus replication and potentially reduce transmission. It is likely that for mediating long-lasting protection, respiratory mucosal vaccines may need to elicit 1) durable antibody-secreting plasma cells that generate secretory IgA antibodies and 2) durable memory immune responses to counteract virus replication in the upper airway. A major roadblock for generating effective respiratory mucosal vaccines is that mucosal antibody responses are dogmatically thought to be short-lived.

Although primary respiratory vaccination can elicit local immunity^22,23^, previous studies have demonstrated that systemic priming followed by an intranasal boost effectively maintains systemic immune responses while significantly enhancing mucosal immune responses^19,32^. To maximize the benefits of this heterologous vaccine regimen, our results suggest that incorporating mucosal adjuvants, such as M7 and CpG, into mucosal vaccine formulations is a strategy to promote robust IgA responses in the respiratory mucosa. We also demonstrate that TLR agonists, which are FDA approved for human use for intramuscular administration, can be repurposed as intranasal adjuvants that elicit durable IgA against SARS-like zoonotic viruses and MERS-CoV. This includes TLR9 agonist CpG, which stimulates type I interferon release and Th1 responses, establishing potent CD8+ and memory T cell populations and mucosal immune responses^33–37^.

Our study demonstrates that M7-CpG adjuvantation of protein-based nanoparticle vaccines successfully elicits durable and protective mucosal IgA in both the lower and upper respiratory tract. While previous studies have evaluated the induction of systemic IgG antibody responses from ferritin nanoparticle protein-based vaccines^27,28,38,39^, to date, it remained unknown if nanoparticle protein-based vaccines may have applications in mucosal delivery. Our study demonstrates that respiratory mucosal delivery of protein-based nanoparticle vaccines elicits long-lasting mucosal IgA responses. Collectively, our findings challenge the conventional dogma that mucosal vaccines do not elicit durable immunity. Moreover, our study suggests respiratory mucosal vaccines that utilize protein nanoparticles coupled with the M7-CpG adjuvant combination is a strategy that elicits long-lasting mucosal IgA, which can reduce the transmission of endemic and emerging zoonotic respiratory coronaviruses.

The development of respiratory mucosal vaccines is inherently constrained by technical and immunological hurdles, including overcoming rapid mucus clearance, targeting nasopharyngeal-associated lymphoid tissue (NALT), generating long-lasting memory responses in airway immune-draining lymph nodes, and developing safe mucosal adjuvants devoid of neurotoxicity risk^14^. Clinically translating the mastroparan-7 (M7) adjuvant requires comprehensive safety validation. A potential limitation of our study is that we do not know which types of dendritic cell (DC) subsets in the respiratory mucosa mediate ferritin nanoparticle vaccine uptake or if respiratory mucosal delivery elicits CD4^+^ and/or CD8^+^ T cell responses. To date, no mucosal adjuvants have been clinically approved for intranasal administration. Although bacterial enterotoxins, such as cholera toxin, are considered the gold standard mucosal adjuvants for effectively driving IgA class switching and Th17 responses, their clinical application in nasal influenza vaccines led to Bell’s palsy^40^. Evaluating whether it triggers excess neuroinflammation or pathological alterations in nasal turbinate histopathology is required in future studies.

Altogether, our results demonstrate that eliciting durable IgA responses in the respiratory mucosa via vaccines is achievable. The requirement of IgA-mediated protection in the respiratory airway from coronavirus infection has important implications for the design of vaccines or countermeasures against emerging, re-emerging, and zoonotic coronaviruses that aim to optimally protect in the respiratory tract.

## Materials and Methods

All coronavirus mouse infections were performed in an animal biosafety level 3 (ABSL3) facility with approval from the Yale Institutional Animal Care and Use Committee and Yale Environmental Health and Safety.

### Study design

The objective of this study was to provide proof of concept in the preclinical development of a mucosal vaccine strategy, demonstrating that an intranasal booster with a mucosal adjuvant could mount durable mucosal IgA responses and breadth of protection against betacoronaviruses.

### Animals and challenge viruses

Eight-week-old female specific-pathogen-free (SPF) hemizygous K18-hACE2 transgenic mice (strain: B6.Cg-Tg(K18-ACE2)2Prlmn/J) were purchased from The Jackson Laboratory (Bar Harbor, ME, USA) and bred in-house. For the viral challenge, all vaccinated mice were anesthetized by using 30% isoflurane diluted in propylene glycol and infected intranasally with 1×10^5^ plaque-forming units (PFU) (50 µL) of RsSHC014, Pangolin-CoV, and SARS-CoV-2 MA10 (BEI resources). Mice with an altered DPP4 at amino acid positions 288 and 330 were bred in-house, and MERS-CoV challenge studies were done with mouse-adapted MERS-CoV m35c4 as described previously^41^. At 2 days post-infection (dpi), the lungs, nasal turbinate, nasal wash, BALF, and serum were collected from each vaccination group (*n* = 5 animals per group).

### Vaccine design and adjuvant formulation

SarbConS, a SARS-like virus synthetic spike protein, was designed by creating a consensus sequence of 23 SARS-like viruses of zoonotic origin. Subsequently, SarbConS was fused to ferritin nanoparticles (FNs) to form a multivalent display as described previously ^39^. The ferritin nanoparticles and SarbConS protein were produced and purified to be endotoxin free. M7-CpG nanoparticles (NPs) were prepared by fabricating M7 peptide (10 nmol = 14.22 µg) (Biomatic, Wilmington, DE, USA) and CpG ODN 1826 (Invivogen, San Diego, CA, USA) (10 µg for N/P = 1) at optimized N/P ratios, followed by sonication at 100% amplitude for 20 minutes and at room temperature (RT)^25^.

### Vaccination

To achieve enhanced mucosal immune responses elicited by vaccination, K18-hACE2 female mice (8-10 weeks old; Jackson Laboratory) bred in-house were immunized using a heterologous intramuscular prime, intranasal boost, followed by a final intranasal boost regimen. Briefly, baseline sera and feces were collected before the priming. Mice were then primed intramuscularly with 1 µg of SARS-CoV-2 mRNA-LNP (Comirnaty (BNT162b2) mRNA-LNP, Pfizer-BioNTech) in a total volume of 50 µL. At 3 and 6 weeks post-prime, the animals received two consecutive intranasal boosts containing 3 µg of SarbConS ferritin nanoparticles, either with or without adjuvants. Mice were immunized intranasally with 3 µg of SarbConS FNs admixed with M7-CpG nanoparticles, administered at a volume of 7.5 µL into each nostril. Sera and fecal samples were collected 2 weeks after the prime and each boost, as well as at the specific time points indicated in each experimental design. Bronchoalveolar lavage fluid (BALF), nasal washes, lungs, nasal turbinates, and serum were collected as part of a terminal collection.

### Enzyme-linked immunosorbent assays (ELISAs)

ELISAs were conducted in 384-well ELISA plates (Greiner Bio-One #781061) coated with 0.5 µg/mL recombinant SARS-CoV-2 spike protein (Sino Biological #40589-V08B1) in cold 1X phosphate-buffered saline (PBS) with 0.1% Tween-20 overnight at 4 °C. For measuring IgA titer against zoonotic bat SARS-like viruses, plates were coated with 1 µg/mL of BatCoV RaTG13 spike, BANAL-236 S2P, BatCoV RsSHC014 RBD, and Pangolin-CoV GXP4L RBD. Plates were washed with 1X Tris-Buffered Saline (TBS) with 25 mM Tris, 0.15 M NaCl, and 0.05% Tween-20 at pH 7.5 and blocked with 1X PBS with 0.1% Tween-20 containing 3% skim milk for 1 hour at room temperature. Serum, nasal wash, and BALF were diluted in dilution solution (PBS-T containing 3% skim milk). To assess mucosal IgA titer as a non-invasive method, fecal samples were prepared for ELISA by homogenizing in 10% goat serum (MP Biomedical, Santa Ana, CA, USA) in 1X PBS and collecting the supernatant after centrifugation, as previously described^25^. Nasal washes were collected in cold 1X PBS and briefly centrifuged to obtain a clear supernatant. Samples were incubated on plates in three-fold serial dilutions overnight at 4 °C. Plates were washed six times with TBS-T by using a plate washer, and 30 µL of anti-mouse IgG (Southern Biotech #1033-01), anti-mouse IgA (Southern Biotech #1040-01), and HRP anti-goat IgG (Southern Biotech #6163-05) was added to each well. After 1 hour of incubation at room temperature or overnight at 4 °C, plates were washed six times with TBS-T. 50 µL of TMB substrate reagent (Sera care #5120-0083) was added to plates at room temperature for 10 min (IgG) or 30 min (IgA), then 50 µL of stop solution (Sera care #5150-0021) was added to terminate the reaction. The absorbance was measured at 450 nm. All incubation steps in the 384-well ELISA were preceded by a brief centrifugation step to ensure complete well wetting and mixture homogeneity. GraphPad Prism was used to determine the area under the curve (AUC) from binding of serial dilutions.

### Flow cytometry

Mice were euthanized with 100% isoflurane, and the lung, mediastinal lymph nodes, and cervical lymph nodes were collected for antigen-specific memory B cell quantitation. For single-cell preparation, lungs were digested in a digestion cocktail containing collagenase A (Roche) and DNase I (Sigma-Aldrich) in RPMI at 37 °C for 30 minutes, followed by dissociation through a 70-µm cell strainer. Mediastinal or cervical lymph nodes were mechanically disrupted using a 1-mL syringe plunger and passed through a 70-µm cell strainer. The isolated cells were treated with ACK buffer for 2 minutes for red blood cell lysis and washed with 1X PBS. The cell suspensions were then stained with Fixable Zombie NIR cell viability dye (BioLegend #423105, 1:200) and incubated with anti-mouse CD16/CD32 Fc Block (BioLegend #101302, 1:100) at 4°C for 30 minutes and subsequently washed with FACS buffer (BioLegend #420201) before cell surface staining. For antigen-specific B cell analysis, SARS-CoV-2 spike or SarbConS spike protein was biotinylated using EZ-Link Sulfo-NHS-LC-Biotin (ThermoFisher #A39257) according to the manufacturer’s instructions. Subsequently, SARS-CoV-2 spike or SarbConS tetramers were prepared via incubation of the biotinylated proteins with streptavidin-PE (Prozyme #PJRS25) or streptavidin-APC (Prozyme #PJ27S) as a stepwise binding at a 4:1 molar ratio at 4 °C. Cells were stained with PE-SarbConS or SARS-CoV-2 spike tetramer and APC-SarbConS or SARS-CoV-2 spike tetramer for 1 hour at 4°C, then further stained with anti-CD3ε (BioLegend #100215, 1:100), anti-B220 (BioLegend #103235, 1:200), anti-CD19 (BioLegend #115537, 1:50), anti-CD38 (BioLegend #102719, 1:200). Cells were washed with FACS buffer, followed by fixation buffer for 20 minutes at 4 °C. After centrifuging at 600 g for 5 minutes, the supernatant was discarded, and cells were washed three times in 1X Perm/Wash buffer using BD Cytofix/Cytoperm Fixation/Permeabilization kit (BD bioscience #554714). Intracellular staining was performed with anti-IgD (BioLegend #405721, 1:200) for 30 minutes at 4°C. Flow cytometry data were acquired on a BD LSR Fortessa Flow Cytometer, and BD FACS Diva software (v.9.3.1) and FlowJo (v.10.10.0) were used for analysis. The gating strategy is provided in Fig S3.

### Pseudovirus neutralizing antibody assay

Neutralizing activity of mouse serum, nasal wash, and bronchoalveolar lavage fluid (BALF) was determined using vesicular stomatitis virus (VSV) pseudovirus neutralization assay as previously described^42^. VSV pseudoviruses expressing Renilla luciferase and deficient in native VSV-G, coated with either SARS-CoV-2 Spike (WA01) or BANAL-236 Spike were produced in 293T cells per previously established protocol^43,44^. VeroE6 cells stably expressing human ACE2 and TMPRSS2 (VeroE6-AT, kindly provided by Dr. Barney Graham) were seeded at 5 x 10^3^ cells/well in 384-well plates (Greiner) in 25 µL of D10/1/AG (DMEM supplemented with 10% FBS, 1% PS, 0.5 µg/mL Amphotericin 0.5 µg/mL and 100 µg/mL G418) prior to infection. Dilutions of sera, nasal wash, and BALF were performed in D0/1/AG (DMEM supplemented with 1% PS, 0.5 µg/mL Amphotericin 0.5 µg/mL and 100 µg/mL G418). Pseudovirus was mixed 1:4 (v/v) with eight 2-fold serial dilutions of serum starting at a 1:20 dilution and incubated for 1 hour at room temperature (5 µL virus + 20 µL serum dilution). For nasal wash and BALF, initial sample volumes were limiting, hence the virus was mixed 1:4 (v/v) with a 1:10 dilution of nasal wash or BALF before incubation for 1 hour at room temperature (5 µL virus + 20 µL of 1:10 nasal/BALF dilution). After a 1-hour incubation, 25 µL of the virus-containing mixture was transferred to the cell-containing wells and incubated at 37 °C with 5% CO_2_ for 24 hours. Following this incubation, Renilla Luciferase Assay System (Promega) was employed as per manufacturer’s protocol. Briefly, supernatant was completely removed from the cells, and 10 µL of Renilla Luciferase Assay Lysis Buffer was added to each well, and incubated for 30 minutes at room temperature on a shaker. After cell lysis, 10 µL of Renilla Luciferase Assay Reagent was added to the wells, and activity was measured on a Cytation5 (BioTek). All neutralization tests were conducted in technical duplicate. The relative 50% inhibitory concentration (IC_50_) for each sample was calculated by performing a four-parameter logistic regression using GraphPad Prism 11 software. Neutralization assays were performed blinded as to sample condition.

### Analysis of viral titer by plaque assays

The lung (right lobes) or nasal turbinate was homogenized in a bead homogenizer tube Lysing Matrix D (MP Biomedical #116913100), containing 1 mL of PBS. After centrifugation (10 minutes, 3000 g), 10-fold serial dilutions of homogenates were prepared in serum-free DMEM and then absorbed on 12-well plates of Vero E6 ACE2/TMPRSS2 cell monolayers for 1 hour at 37 °C. Nasal washes or oral swabs were performed using polyester tipped applicators (Pruitan PurFlock Ultra 25-3206-U) and were obtained by flushing the nasal cavity or gently swabbing the oral cavity with 500 µL of cold sterile 1X PBS with 1% BSA and stored at −80 °C for further assays. The cell monolayers were then overlaid with DMEM containing 1.2% Avicel. After 42 - 45 hours of incubation at 37 °C, the overlay media was aspirated, and the cells were fixed with 10% formalin. Following fixation, the monolayers were stained with crystal violet and washed with water for plaque visualization.

### Statistical analyses

Statistical significance among groups was evaluated using a two-way ANOVA with Tukey’s post-hoc test for the ELISA assay. For plaque assays and flow cytometry, comparisons were analyzed via the non-parametric Kruskal-Wallis test followed by Dunn’s multiple comparisons test. All data are presented as mean ± SEM. P-values less than 0.05 were considered statistically significant (ns, not significant; *p < 0.05, **p < 0.01, ***p < 0.001, and ****p < 0.0001). P values were adjusted for multiple comparisons. All tests were performed using GraphPad Prism software (version 11).

## Author contributions

Conceptualization: D.R.M.

Methodology: J.S., E.B., B.M., O.M., B.M., and R.F.

Investigation: J.S. E.B., B.M., O.M., B.M., and R.F.,

Visualization: J.S. B.M., and D.R.M.

Funding acquisition: D.R.M., and K.O.S.,

Project administration: D.R.M.

Supervision: D.R.M., C.W., B.I.

Writing – original draft: J.S., and D.R.M.

Writing – review & editing: J.S., E.B., B.M., O.M., B.M., R.F., K.O.S., C.W., B.I., D.R.M.

## Supplementary materials include

Fig. S1-S4.

