## Supplemental Figures for "Mucosal vaccine-elicited IgA is protective against zoonotic Betacoronavirus challenge"

### Supplemental figure legends

**Figure S1**

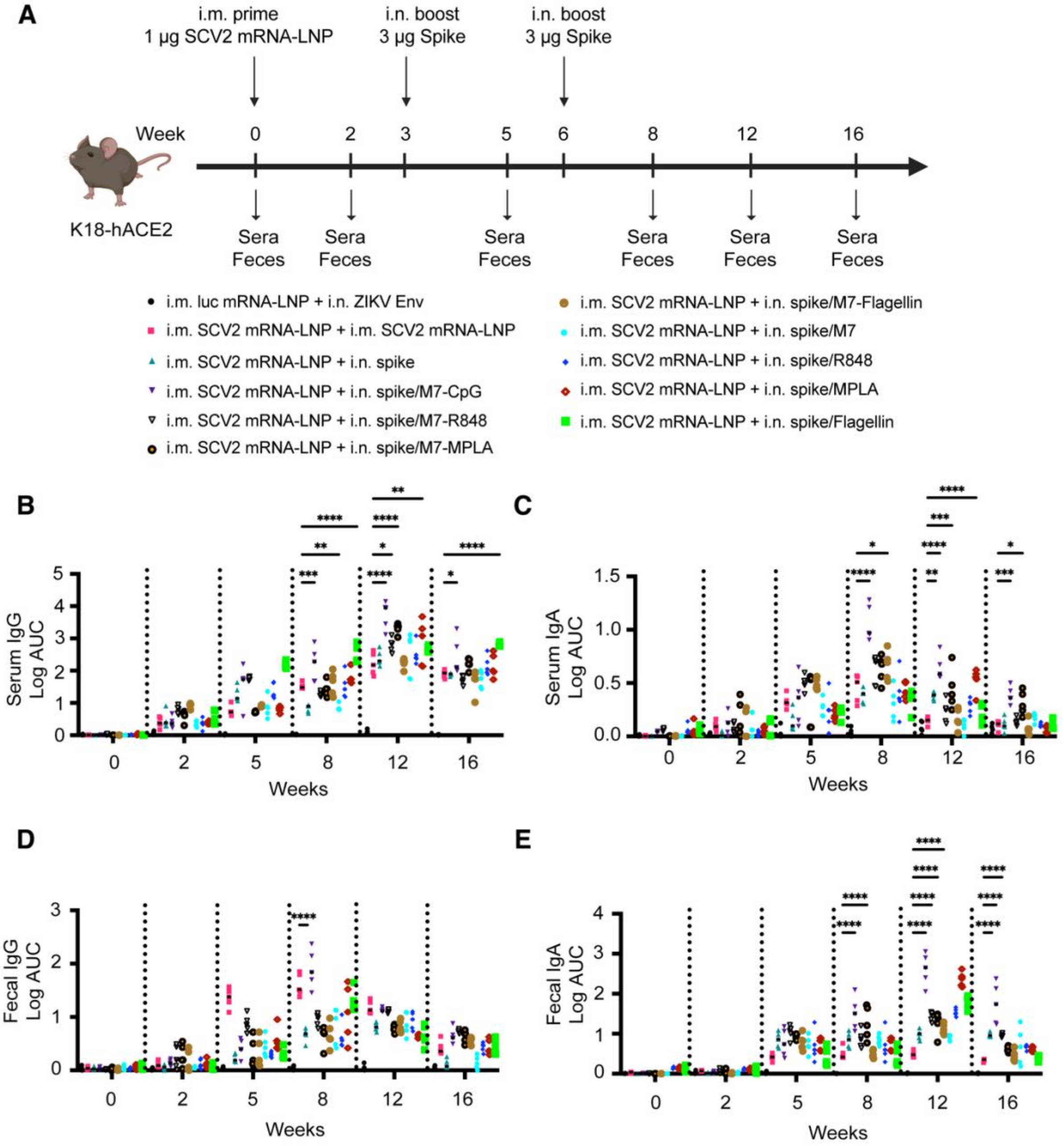

**Figure S1. Self-assembled nanoparticle mucosal adjuvant combination of mast cell agonist mastroparan-7 (M7) and TLR9 agonist CpG elicits highest magnitude and durable mucosal IgA responses.**

Immunogenicity in mice intranasally boosted with SARS-CoV-2 spike protein formulated with various combinations of TLR agonist and the mast cell activator mastoparan-7 (M7) in SARS-CoV-2-immune model mice. **(A)** Schematic of the vaccination strategy. **(B-E)** Measurement of IgG or IgA titers by ELISA. K18-hACE2 mice were immunized intramuscularly at week 0 and intranasally at weeks 3 and 6. Sera and feces were collected 1 day before priming (pre-prime), 2 weeks after priming or boosting, and at the indicated weeks ($n$ = 5 per group). Fecal samples were analyzed as a non-invasive measure to evaluate mucosal IgA responses. Data are presented as mean ± SEM. Statistical significance was determined using two-way ANOVA followed by Tukey's multiple comparisons test. *$p < 0.05$, **$p < 0.01$, ***$p < 0.001$, and ****$p < 0.0001$.

### Figure S2

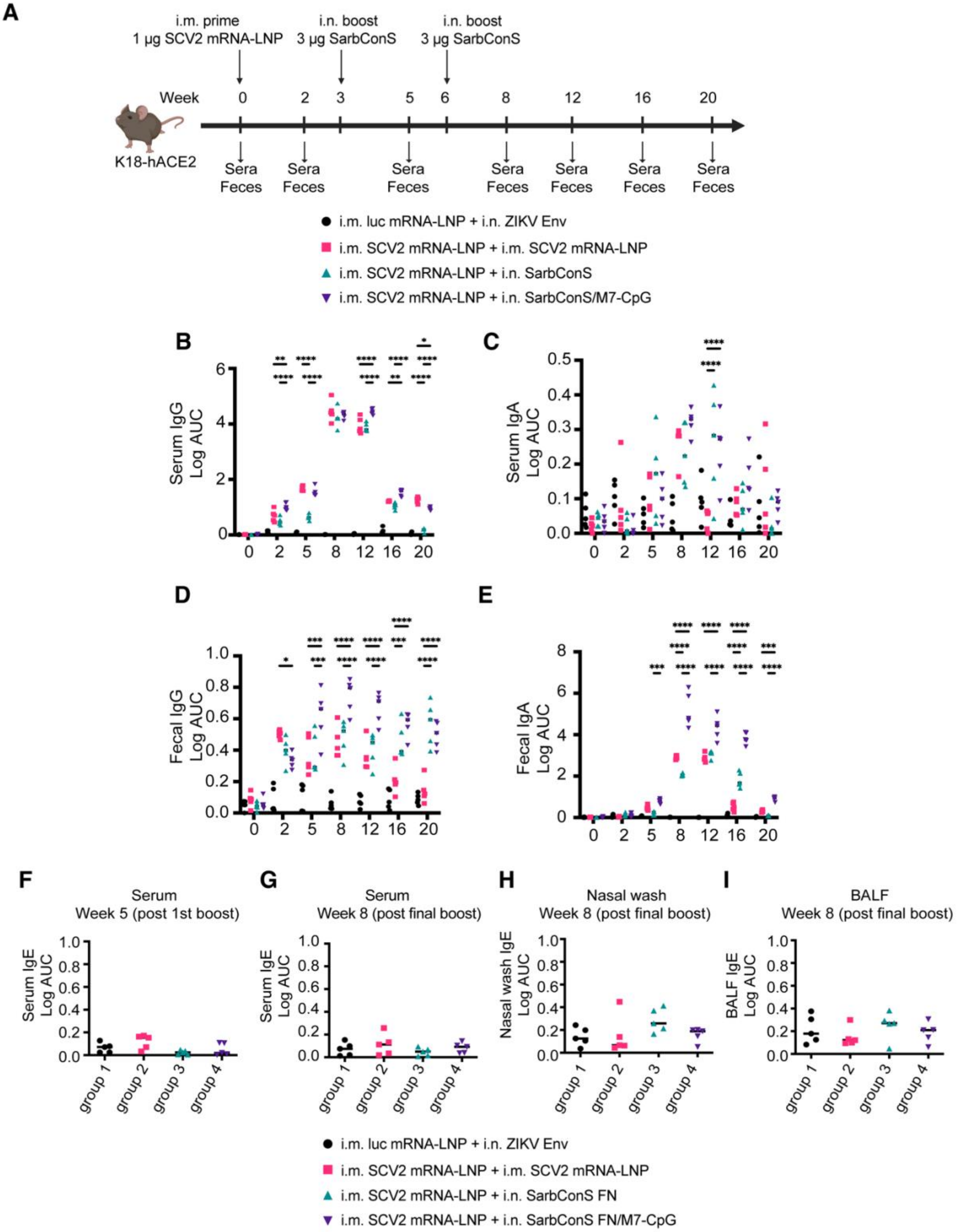
A
i.m. prime
1 µg SCV2 mRNA-LNP
i.n. boost
3 µg SarbConS
i.n. boost
3 µg SarbConS
Week
0 2 3 5 6 8 12 16 20
K18-hACE2
Sera Feces
i.m. luc mRNA-LNP + i.n. ZIKV Env
i.m. SCV2 mRNA-LNP + i.m. SCV2 mRNA-LNP
i.m. SCV2 mRNA-LNP + i.n. SarbConS
i.m. SCV2 mRNA-LNP + i.n. SarbConS/M7-CpG
B
Serum IgG Log AUC
C
Serum IgA Log AUC
D
Fecal IgG Log AUC
E
Fecal IgA Log AUC
F
Serum
Week 5 (post 1st boost)
Serum IgE Log AUC
G
Serum
Week 8 (post final boost)
Serum IgE Log AUC
H
Nasal wash
Week 8 (post final boost)
Nasal wash IgE Log AUC
I
BALF
Week 8 (post final boost)
BALF IgE Log AUC
group 1
group 2
group 3
group 4
i.m. luc mRNA-LNP + i.n. ZIKV Env
i.m. SCV2 mRNA-LNP + i.m. SCV2 mRNA-LNP
i.m. SCV2 mRNA-LNP + i.n. SarbConS FN
i.m. SCV2 mRNA-LNP + i.n. SarbConS FN/M7-CpG

**Figure S2. An intranasal nanoparticle booster admixed with M7-CpG outperforms an intramuscular mRNA-LNP booster in eliciting durable mucosal IgA responses.** Assessment of spike-specific mucosal IgA and IgG titers following intranasal SarbConS versus intramuscular mRNA-LNP boosting. **(A)** Experimental timeline of the vaccination and specimen collection. **(B-E)** Serum and fecal IgG or IgA titers were determined by ELISA. K18-hACE2 mice were immunized intramuscularly at week 0 and intranasally at weeks 3 and 6. Sera and feces were collected 1 day before priming (pre-prime), 2 weeks after priming or boosting, and at the indicated weeks for longitudinal analysis (*n* = 5 per group). **(F-I)** spike-specific IgE titers in serum at week 5 (post $1^{st}$ boost) **(F)** and in serum **(G)**, nasal wash **(H)**, and BALF **(I)** at week 8 (post $2^{nd}$ boost). Data are presented as mean ± SEM. Statistical significance was determined using two-way ANOVA followed by Tukey's multiple comparisons test. *$p < 0.05$, **$p < 0.01$, ***$p < 0.001$, and ****$p < 0.0001$.

**Figure S3**

**A**

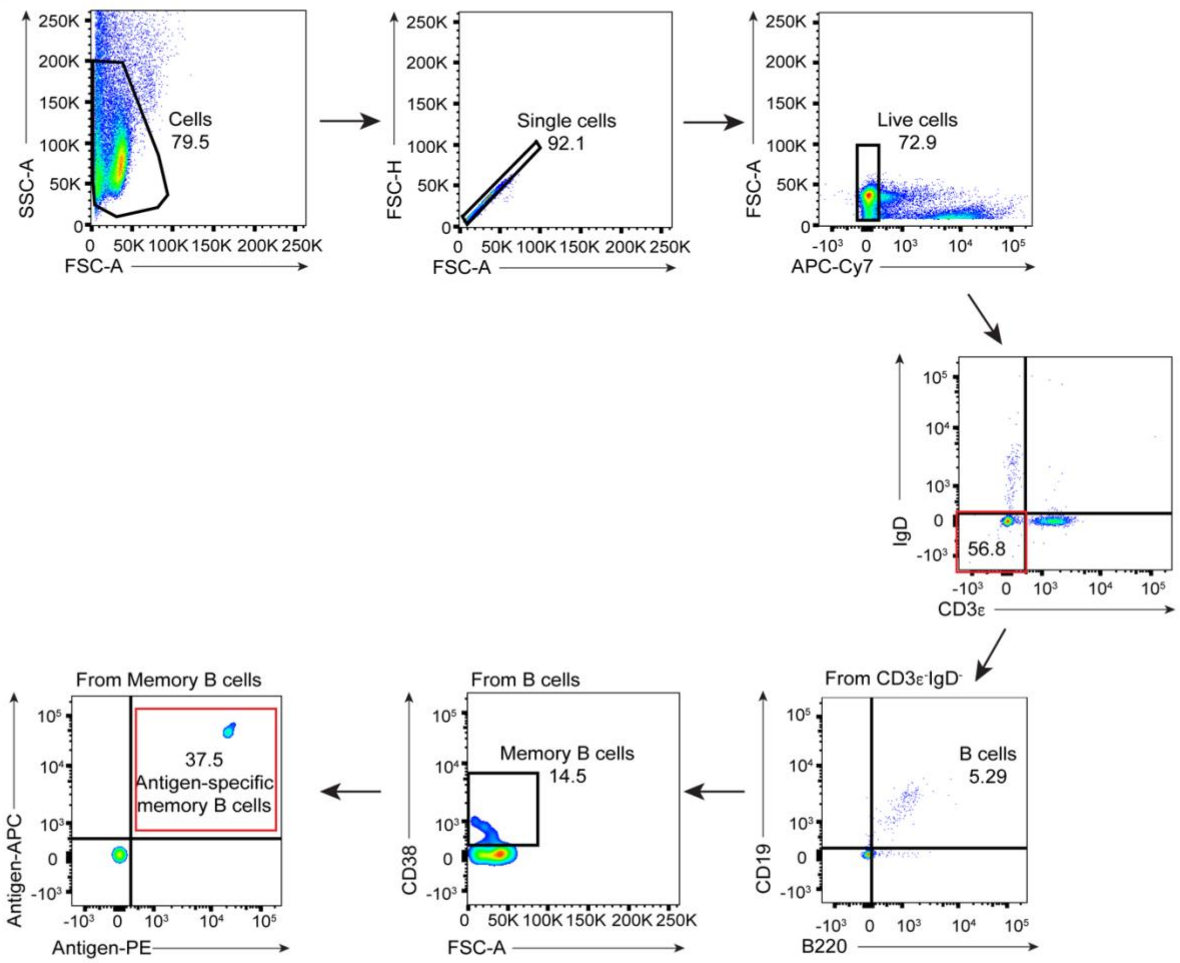


**Figure S3. Gating strategies for antigen-specific memory B cells.**

Gating strategies of antigen-specific CD3ε$^-$IgD$^-$B220$^+$CD19$^+$CD38$^+$ memory B cells from lung, mediastinal and cervical lymph nodes.

**Figure S4**

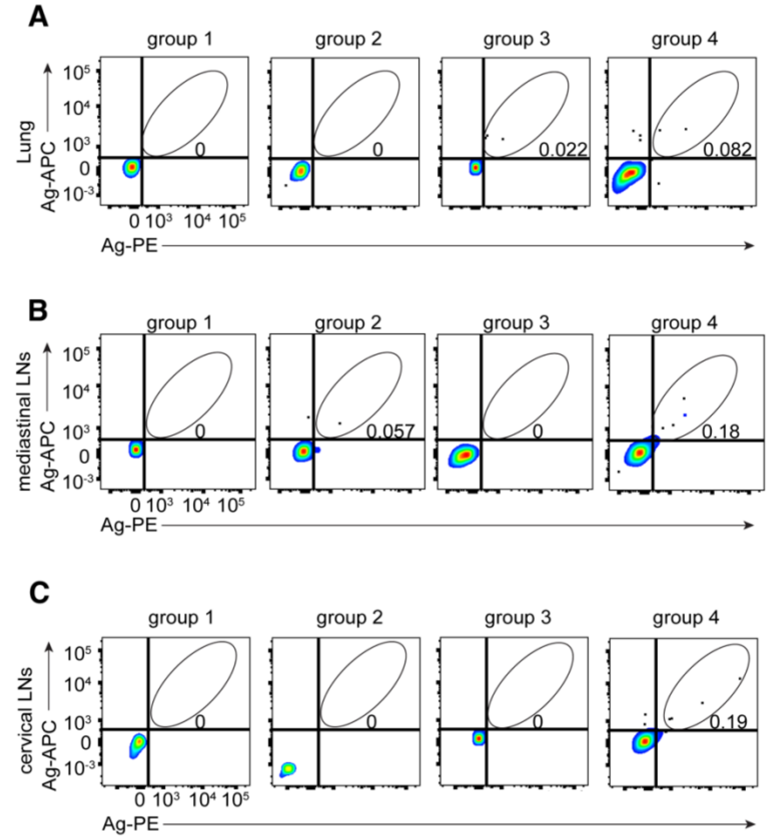


**Figure S4. Intranasal boosting with SarbConS ferritin nanoparticle adjuvanted with M7-CpG elicits durable antigen-specific memory B cell responses detectable at week 24.**

Flow cytometry analysis of antigen-specific memory B cell frequencies in lungs **(A)**, mediastinal lymph nodes **(B)**, and cervical lymph nodes **(C)**. **(A-C)** Representative flow cytometry plots of antigen-specific CD3$\varepsilon^-$IgD$^-$B220$^+$CD19$^+$CD38$^+$ memory B cells at week 24.
